# An oomycete effector that induces shade avoidance like growth and suppresses plant defenses targets the AUX/IAA protein IAA11

**DOI:** 10.1101/2025.02.06.636700

**Authors:** Maria Florencia Bogino, Juan Marcos Lapegna Senz, Nicolás Tamagnone, Lucille Tihomirova Kourdova, Andrés Romanowski, Lennart Wirthmueller, Georgina Fabro

## Abstract

Filamentous pathogens, such as oomycetes, employ multiple strategies to suppress plant immunity, one of which involves manipulating phytohormone signaling pathways. The effector HaRxL106 from Hyaloperonospora arabidopsidis (Hpa), known to induce shade avoidance syndrome (SAS)-like growth, has previously been shown to interact with brassinosteroid signaling components. Here, we investigate the role of HaRxL106 in altering auxin signaling, revealing that HaRxL106 targets the AUX/IAA protein IAA11, a key negative regulator of auxin responses. Our data suggest that HaRxL106 impedes the repressive function of IAA11, enhancing auxin signaling and promoting SAS, which concurrently suppresses plant immune responses. These findings provide insights into how pathogen effectors manipulate plant growth and immunity to facilitate infection.

## Main manuscript

Filamentous pathogens, like oomycetes, exhibit multiple mechanisms to subdue plant immunity (Fabro, 2022). Altering phytohormone pathways that regulate plant growth is one of them (Han and Kahmann, 2019). For example, the oomycete *Hyaloperonospora arabidopsidis* (Hpa) that infects *Arabidopsis thaliana*, produces the effector HaRxL106, which interacts with the transcription factor BIM1 (BES1-INTERACTING MYC-LIKE 1), affecting brassinosteroid (BR) signaling (Bogino *et al*., 2024). HaRxL106 expression in Arabidopsis causes shade avoidance syndrome (SAS)-like phenotypes (Wirthmueller et al., 2018). In plants displaying SAS, BR and auxin signaling are activated (Casal, 2013; Keuskamp *et al*., 2011) while immune responses are compromised (De Wit *et al*., 2013). Here we report that HaRxL106 also targets IAA11, a protein involved in auxin signaling, revealing new insights into this effector’s function.

As the activation of the auxin pathway is required for SAS establishment, we wondered if Hpa infection might induce auxin signaling. Previous data (Wang *et al*., 2007) suggested that the auxin reporter DR5:GUS was induced in Hpa infected tissues. We repeated this assay quantifying the activation of the reporter at different times post-infection with Hpa isolate NoCo2 (Supplemental Methods 1, 2 and 6). We observed a statistically significant increase of GUS staining in infected cotyledons at 3 and 7 days post-inoculation (Figures 1A, 1B). To investigate if Hpa effector HaRxL106 was responsible for this, we used the pEDV system to deliver it into DR5:GFP plants (Fabro *et al*., 2011; Supplemental Methods 3). We observed that pEDV6-HaRxL106 increased GFP fluorescence, but the variability of this transient assay affected statistical significance (Supplemental Figure 1). Therefore, we generated stable transformants of DR5:GUS plants carrying an estradiol-inducible HaRxL106 construct (pER8:HA-HaRxL106) (Supplemental Methods 1). We detected a statistically significant increase in GUS staining in these seedlings upon estradiol treatment (Figure 1C).

**Figure 1:**
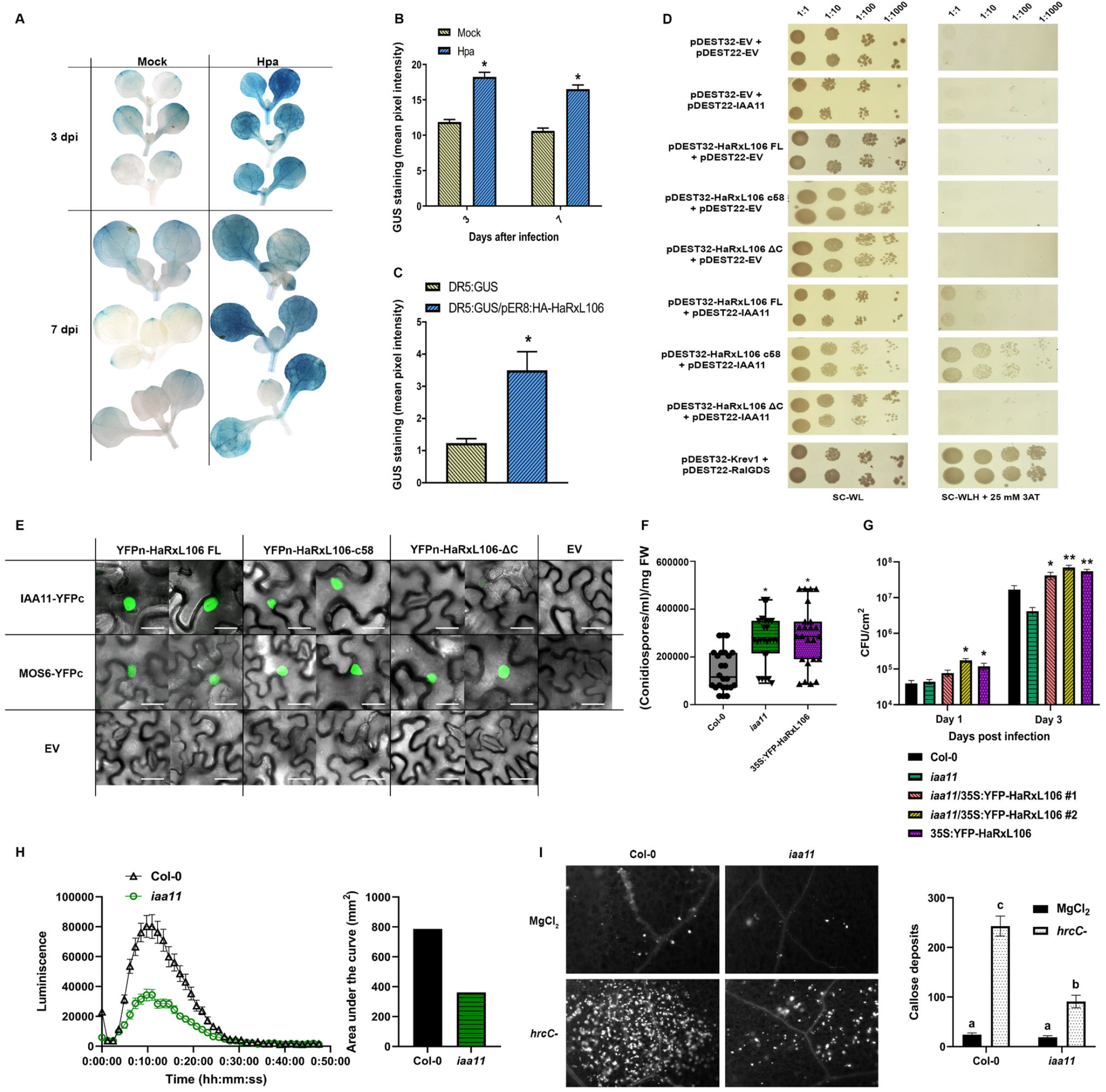
The oomycete effector HaRxL106 induces auxin signaling and its target IAA11 positively affects plant defenses. **A**. Representative images of seedlings expressing the DR5:GUS construct either uninfected (mock) or 3 and 7 days post-infection with Hpa NoCo2. **B**. Quantification of the GUS staining intensity of the cotyledons of seedlings (n=75) like those shown in A. Asterisks indicate significant differences between treatments with p<0.001 according to a non-parametric ANOVA (Kruskal-Wallis). **C**. Quantification of GUS staining intensity in DR5:GUS and DR5:GUS/pER8:HA-HaRxL106 10-day old seedlings (n=24) treated with 1.25 μM β-Estradiol. Asterisks indicate significant differences between treatments with p<0.001 according to a non-parametric ANOVA (Kruskal-Wallis). **D**. Y2H assay using the effector full length sequence (HaRxL106 FL) and its carboxy-terminal (HaRXL106-c58) and amino-terminal (HaRxL106-ΔC) domains. SC-WL: Synthetic Complete medium for yeast lacking Tryptophan and Leucine. SC-WLH + 25 mM 3-AT: Synthetic Complete medium for yeast lacking Tryptophan, Leucine, and Histidine, supplemented with 25 mM 3-Aminotriazole. pDEST32-Krev1 + pDEST22-RalGDS: positive interaction control. pDEST32: bait vector. pDEST22: prey vector. **E**. Representative confocal microscopy images of the abaxial epidermis of *N. benthamiana* leaves expressing the indicated constructs. MOS6-YFPc + YFPn-HaRxL106: positive interaction control (Wirthmueller *et al*., 2015). As negative controls, each construct was expressed individually. Scale bars represent 20 μm. **F**. Amount of conidiospores produced by Hpa on *A. thaliana* seedlings of the specified genotypes. Significant differences between Col-0 and the other genotypes based on a non-parametric ANOVA (Kruskal Wallis) with p<0.05 are highlighted by asterisks. This experiment was repeated three times with similar results. **G**. *Pseudomonas syringae pv. tomato* (Pst) DC3000 colony-forming units per square centimeter of leaf (CFU/cm^2^) determined on the mentioned genotypes at 1 and 3 days post-inoculation. Significant differences compared to the wild-type genotype (Col-0) are indicated according to the t-test for mean differences with p<0.05 (*) and p<0.01 (**), n=6. **H**. Kinetics of the oxidative burst in the apoplast (apoplastic ROS) following treatment with 100 nM flg22 (left panel). The right panel shows the quantitation of the area under the curves shown in the left panel. **I**. Representative images of the abaxial epidermis of leaves from the indicated genotypes treated with the vehicle (10 mM MgCl_2_) or inoculated with Pst *hrcC*-mutant (left panel). Assessment of callose deposits in the photographed fields (right panel). Different letters indicate statistically significant differences with p<0.05 according to a non-parametric ANOVA (Kruskal Wallis), n=36.

Next, we took advantage of a yeast-two-hybrid (Y2H) screen performed with HaRxL106 (Dr. Jonathan D.G. Jones, personal communication) and a library of ~8000 Arabidopsis proteins, which suggested that HaRxL106 directly interacts with IAA11 (INDOLE-3-ACETIC ACID INDUCIBLE 11). IAA11 is a canonical AUX/IAA that negatively regulates auxin responses (Leyser, 2018) and represses root growth (Mielecki *et al*., 2022). IAA11 interacts with several ARFs (Auxin Response Factors) (Piya *et al*., 2014) and with another HaRxL106 target, RCD1 (Jaspers *et al*., 2009). To verify HaRxL106-IAA11 interaction, we performed a Y2H assay using different domains of HaRxL106: full-length (FL), C-terminal (c58), and amino-terminal (ΔC) (Figure 1D; Supplemental Methods 4). Yeast co-expressing HaRxL106-c58 and IAA11, and to a lesser extent those containing HaRxL106 FL and IAA11, grew on selective media, indicating that mainly the C-terminal domain of HaRxL106 mediates the interaction with IAA11. To confirm if the interaction occurred *in planta*, we performed Bi-molecular fluorescence complementation-BiFC-assays in *Nicotiana benthamiana* (Supplemental Methods 5). We found that HaRxL106-FL and HaRxL106-c58, interacted with IAA11 in the plant cell nucleus (Figure 1E).

To determine if IAA11 was relevant for Hpa infection, we phenotyped the *iaa11* mutant (SALK_033787), previously reported to be more susceptible to Hpa race Emwa1 (Wessling *et al*., 2014). This line is a knock-down for the *IAA11* transcript (*iaa11-4* in Mielecki et al., 2022, Supplemental Methods 1; Supplemental Figure 2). Notably, the compatible Hpa Noco2 isolate sporulated to a higher level on *iaa11* mutants, similar to the elevated susceptibility observed for HaRxL106 over-expressors (35S:YFP-HaRxL106) (Figure 1F). Conversely, *iaa11* mutants were not more susceptible to the hemibiotrophic bacterium *Pseudomonas syringae* (Pst) (Figure 1G). Consistently, two hallmarks of PAMP Triggered Immune (PTI) responses; flg22-triggered ROS production (Figure 1H) and Pst *hrcC*-induced callose deposition (Figure 1I), were diminished in *iaa11* mutants, as previously observed for 35S:YFP-HaRxL106 plants (Fabro *et al*., 2011; Supplemental Methods 7). These results suggest that IAA11 might function as a positive regulator of PTI.

**Figure 2:**
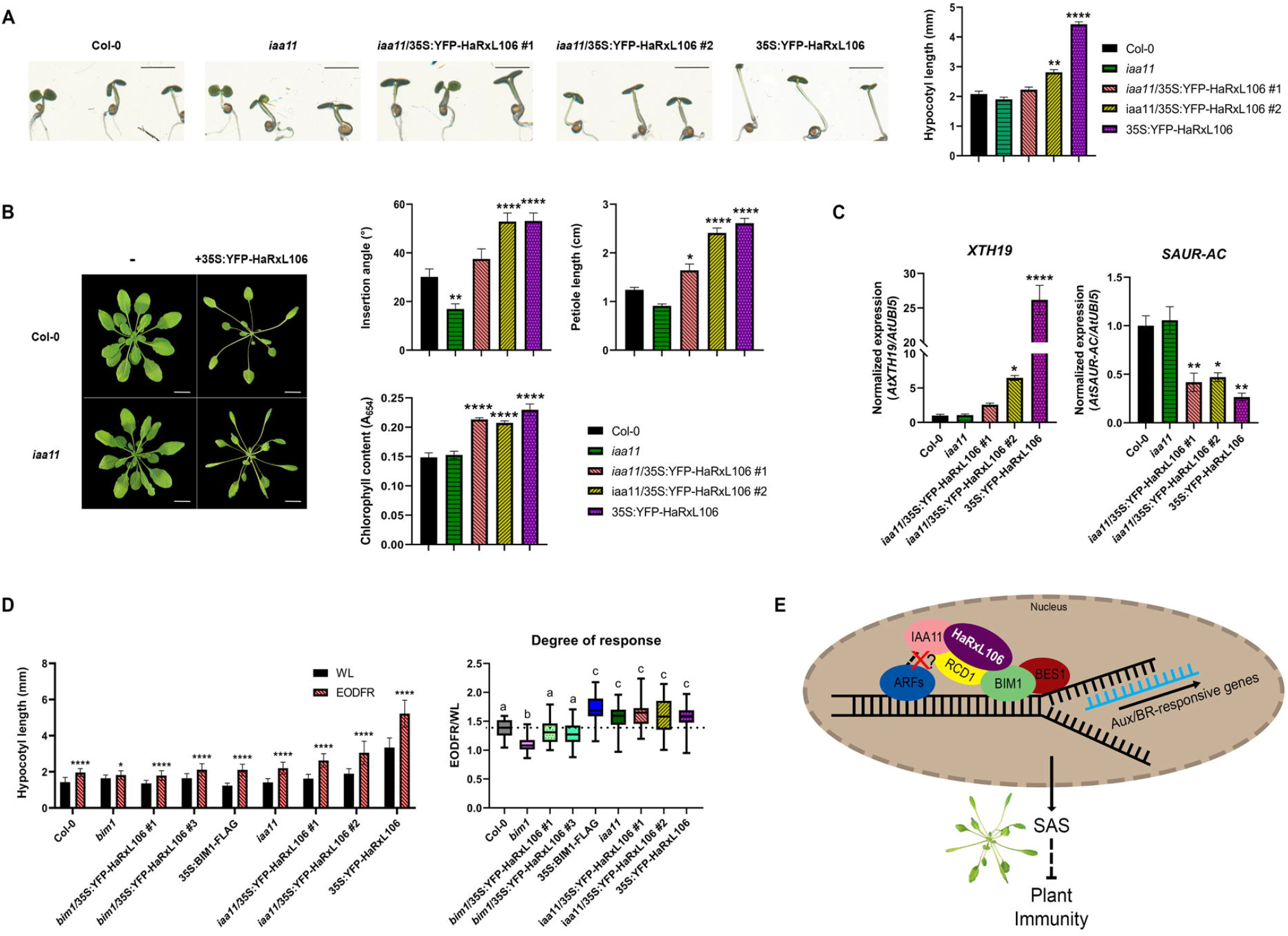
HaRxL106 impedes IAA11 negative regulatory functions over auxin signaling. **A**. Representative pictures of 7-day-old seedlings of the specified genotypes (left panel) used to quantitate their hypocotyl length (graph in right panel). Scale bars: 2 mm. Significant differences between genotypes and the control (Col-0) are indicated by asterisks (**: p<0.01; ****: p<0.0001) according to a non-parametric ANOVA (Kruskal Wallis), n = 26. **B**. Images depicting representative two-month-old plants that overexpress HaRxL106 in either wild type (Col-0) or *iaa11* mutant background (Col-0/35S:YFP-HaRxL106 and *iaa11*/35S:YFP-HaRxL106 #2, left panel). Scale bars represent 2 cm. Right panel shows the quantification of petiole length, petiole insertion angle, and chlorophyll content. Significant differences between the genotypes and the control plants (Col-0) are indicated by asterisks (*: p<0.05; **: p<0.01; ****: p<0.0001) according one-way ANOVA with Tukey’s multiple comparisons test. **C**.Expression of the indicated genes normalized to *AtUBI5* (reference gene) in 7-day-old seedlings of the indicated genotypes. Asterisks denote statistically significant differences between the genotypes and Col-0 (*: p<0.05; **: p<0.01; ****: p<0.0001) according to a one-factor ANOVA (Tukey post hoc test). **D**. Left panel shows hypocotyl length of 7-day-old Arabidopsis seedlings of the indicated genotypes, grown in 12 h white light:12 h darkness (WL, control) or 12 h white light + 10’ far red at the end of the day: 11 h 50’ darkness (EODFR). Statistical differences according to a nonparametric ANOVA (Kruskall Wallis) are indicated with asterisks (*: p<0.05 and ****: p<0.0001). In the right panel, the degree of response of the plants shown in the left panel is calculated (EODFR/WL ratio). Different letters denote statistical differences according to a non-parametric ANOVA (Kruskall Wallis) with p<0.05. **E**.Diagram summarizing the effects of HaRxL106 over Auxin and Brassinosteroids signaling pathways and its consequences on plant immunity. In the nucleus of plant cells, HaRxL106 may recruit the repressor protein IAA11, preventing its interaction with ARFs. ARFs would then promote the transcription of auxin responsive genes. Additionally, HaRxL106 interacts with BIM1 possibly stabilizing the BIM1/BES1 heterodimer, activating the transcription of Brassinosteroid response genes. Another known target of HaRxL106, RCD1, which has been reported to interact with IAA11 (Jaspers *et al*., 2009), may act as a scaffold or a co-repressor of IAA11 together with HaRxL106. The coordinated activation of both Auxin and BR signaling triggers a constitutive SAS-like response, which likely has repressive effects on defense responses.

35S:YFP-HaRxL106 plants exhibit constitutive SAS-like growth. This phenotype was largely maintained when the effector was introduced into the *iaa11* background (Figure 2A and B; Supplemental Figure 3; Supplemental Methods 1, 8 and 9). *iaa11*/35S:YFP-HaRxL106 plants had longer hypocotyls and petioles, higher leaf insertion angles and increased chlorophyll content than Col-0. The *iaa11*/35S:YFP-HaRxL106 lines also showed enhanced susceptibility to Pst (Figure 1G). These results suggest that HaRxL106 affects IAA11, disrupting its role as a negative regulator of auxin responses (Leyser, 2018). If so, 35S:YFP-HaRxL106 plants should show altered expression of auxin/shade-responsive genes (*XTH19* and *SAUR-AC-SAUR15-*; Nemhauser et al., 2006; Roig-Villanova et al., 2007). To investigate this, we measured transcript levels (Figure 2C; Supplemental Methods 10). We found that HaRxL106 expression altered *XTH19* and *SAUR-AC* transcripts levels in both Col-0 and *iaa11* backgrounds.

If HaRxL106 represses the negative regulator IAA11, then both *iaa11* mutants and plants with native IAA11 but overexpressing HaRxL106 should exhibit increased responsiveness to SAS-inducing treatments. To investigate this, we grew *iaa11* and 35S:YFP-HaRxL106 seedlings in a shade-mimicking condition (End-of-Day Far-Red, EODFR, Supplemental Methods 9, Mizuno *et al*., 2015; Romanowski *et al*., 2021). As BIM1 is required for SAS, *bim1* mutants were used as negative control and 35S-BIM1-FLAG transgenics as positive control (Cifuentes-Esquivel *et al*., 2013). EODFR treatment induced a higher degree of SAS in *iaa11*, 35S:YFP-HaRxL106 and both *iaa11*/35S:YFP-HaRxL106 lines (Figure 2D). These plants’ degree of response was statistically indistinguishable from BIM1-FLAG plants. These data suggest that the HaRxL106 effector negatively regulates IAA11, impeding its inhibitory function over auxin signaling. Further experiments are needed to determine if HaRxL106 alters IAA11 turnover or impedes its association with ARFs.

A working model summarizing HaRxL106 functions is shown in Figure 2E. We propose that the de-repression of auxin signaling, combined with the induction of BR-signaling, may form a reinforcement loop that triggers a constitutive state of SAS, which in turn negatively affects plant immune responses. Our findings provide additional evidence for how phytopathogen effectors promote infection by modulating plant growth at the expense of defense responses.

## Supporting information

Supplemental Data: Suppl. Figures, Materials and Methods, Tables and References

## Acknowledgments

To NASC/ABRC for *iaa11* seeds. To Dr. Hongtao Liu for *bim1* and BIM1-FLAG seeds. To Dr. Dae Sung Kim and Dr. Jonathan D.G. Jones laboratory (TSL, Norwich, UK) for sharing unpublished data. To Prof. Dr. Tina Romeis for supporting the research stays of GF and MFB at Free University of Berlin and IPB Halle. To Dr. Maria Elena Alvarez for the support to GF and the initial PhD fellowship of MFB To Dr. Nicolas Cecchini for his critical and comprehensive reading of the manuscript. To Dr. Maria Florencia Nota and Dr. Ana Paula Cislaghi for valuable suggestions on experimental work. To Drs. Cecilia Sampedro and Carlos Mas from CEMINCO for their assistance with confocal microscopy. To undergrad students Gonzalo Sánchez and Marco Stefano Donadio Grangetto for their contribution in samples preparation. The authors declare no conflicts of interest.

## Author contributions

GF and MFB conceived research and designed the experiments. MFB generated transgenics and performed most of the experiments. JMLS, NT and LTK collaborated in experiments (Y2H assays, Pst growth curves and qPCRs). AR collaborated with the EOFDR assays and hosted MFB for a stay in his lab. LW hosted MFB while performing experiments with Hpa in his lab. MFB, LW, AR and GF analyzed the data. MFB and GF wrote the manuscript draft. GF and LW secured funding for research. MFB, JMLS, NT, LTK, AR, LW and GF collaborated to generate the final manuscript.

## Supplemental data

(see attached PDF file).

## Funding

Work in GF lab was supported by the Consejo Nacional de Investigaciones Científicas y Técnicas (CONICET), Agencia Nacional de Promoción Científica y Tecnológica (ANPCyT, FONCyT PICT-2017-0515 and PICT-2020-0763), and the Secretary of Science and Technology of Universidad Nacional de Córdoba (SECyT-UNC). GF is a Career Researcher of CONICET-UNC. This work was supported by an Alexander von Humboldt Georg Forster Fellowship (to GF). MFB was a FONCyT-ANPCyT Fellow (project PICT-2018-4588 of Dr. M. E. Alvarez) and was awarded a One-year PhD Grant fellowship by the DAAD to visit Dr. L.W. lab. Also was recipient of a The Company of Biologists Travel Grant to do a research stay at Dr. A.R. lab. MFB was a recipient of a CONICET FinDoc Fellowship. JMLS was recipient of a CIN (Consejo Interuniversitario Nacional) student fellowship at GF lab. Dr. LTK is a postdoc fellow in the GF lab supported by a FONCyT-ANPCyT fellowship, under PICT-2019-02331 and a CONICET fellowship. NT is a biotech undergrad student at Dr. GF lab. AR is PI at Wageningen University and Research, The Netherlands. LW acknowledges core funding from Free University Berlin and the Leibniz Institute of Plant Biochemistry.

## Data availability

All relevant data can be found within the manuscript and its supporting materials.

## Notes

### Competing Interest Statement

The authors have declared no competing interest.

## References

Bogino, M. F., Lapegna Senz, J. M., Kourdova, L. T., Tamagnone, N., Romanowski, A., Wirthmueller, L., & Fabro, G. (2024). Downy mildew effector HaRxL106 interacts with the transcription factor BIM1 altering plant growth, BR signaling and susceptibility to pathogens. The Plant Journal.

Casal, J. J. (2013). Photoreceptor signaling networks in plant responses to shade. Annual review of plant biology, 64(1), 403–427.

Cifuentes-Esquivel, N., Bou-Torrent, J., Galstyan, A., Gallemí, M., Sessa, G., Salla Martret, M., Roig-Villanova, I., Ruberti, I., & Martínez-García, J. F. (2013). The bHLH proteins BEE and BIM positively modulate the shade avoidance syndrome in Arabidopsis seedlings. The Plant Journal, 75(6), 989–1002.

De Wit, M., Spoel, S. H., Sanchez-Perez, G. F., Gommers, C. M., Pieterse, C. M., Voesenek, L. A., & Pierik, R. (2013). Perception of low red: far-red ratio compromises both salicylic acid-and jasmonic acid-dependent pathogen defences in Arabidopsis. The Plant Journal, 75(1), 90–103.

Fabro, G., Steinbrenner, J., Coates, M., Ishaque, N., Baxter, L., Studholme, D. J., … & Jones, J. D. (2011). Multiple candidate effectors from the oomycete pathogen Hyaloperonospora arabidopsidis suppress host plant immunity. PLoS pathogens, 7(11), e1002348.

Fabro, G. (2022). Oomycete intracellular effectors: specialised weapons targeting strategic plant processes. New Phytologist, 233(3), 1074–1082.

Han, X., & Kahmann, R. (2019). Manipulation of phytohormone pathways by effectors of filamentous plant pathogens. Frontiers in Plant Science, 10, 822.

Jaspers, P., Blomster, T., Brosche, M., Salojärvi, J., Ahlfors, R., Vainonen, J. P., … & Kangasjärvi, J. (2009). Unequally redundant RCD1 and SRO1 mediate stress and developmental responses and interact with transcription factors. The Plant Journal, 60(2), 268–279.

Keuskamp, D. H., Sasidharan, R., Vos, I., Peeters, A. J., Voesenek, L. A., & Pierik, R. (2011). Blue-light-mediated shade avoidance requires combined auxin and brassinosteroid action in Arabidopsis seedlings. The Plant Journal, 67(2), 208–217.

Leyser, O. (2018). Auxin signaling. Plant physiology, 176(1), 465–479.

Mielecki, J., Gawroński, P., & Karpiński, S. (2022). Aux/IAA11 is required for UV-AB tolerance and auxin sensing in Arabidopsis thaliana. International Journal of Molecular Sciences, 23(21), 13386.

Mizuno, T., Oka, H., Yoshimura, F., Ishida, K., & Yamashino, T. (2015). Insight into the mechanism of end-of-day far-red light (EODFR)-induced shade avoidance responses in Arabidopsis thaliana. Bioscience, biotechnology, and biochemistry, 79(12), 1987–1994.

Nemhauser, J. L., Hong, F., & Chory, J. (2006). Different plant hormones regulate similar processes through largely nonoverlapping transcriptional responses. Cell, 126(3), 467–475.

Piya, S., Shrestha, S. K., Binder, B., Stewart Jr, C. N., & Hewezi, T. (2014). Protein-protein interaction and gene co-expression maps of ARFs and Aux/IAAs in Arabidopsis. Frontiers in plant science, 5, 744.

Roig-Villanova, I., Bou-Torrent, J., Galstyan, A., Carretero-Paulet, L., Portoles, S., Rodríguez-Concepción, M., & Martínez-García, J. F. (2007). Interaction of shade avoidance and auxin responses: a role for two novel atypical bHLH proteins. The EMBO journal, 26(22), 4756–4767.

Romanowski, A., Furniss, J. J., Hussain, E., & Halliday, K. J. (2021). Phytochrome regulates cellular response plasticity and the basic molecular machinery of leaf development. Plant Physiology, 186(2), 1220–1239.

Wang, D., Pajerowska-Mukhtar, K., Culler, A. H., & Dong, X. (2007). Salicylic acid inhibits pathogen growth in plants through repression of the auxin signaling pathway. Current Biology, 17(20), 1784–1790.

Weßling, R., Epple, P., Altmann, S., He, Y., Yang, L., Henz, S. R., … & Braun, P. (2014). Convergent targeting of a common host protein-network by pathogen effectors from three kingdoms of life. Cell host & microbe, 16(3), 364–375.

Wirthmueller, L., Asai, S., Rallapalli, G., Sklenar, J., Fabro, G., Kim, D. S., Lintermann, R., Jaspers, P., Wrzaczek, M., Kangasjärvi, J., MacLean, D., Menke, F. L. H., Banfield, M. J. & Jones, J. D. (2018). Arabidopsis downy mildew effector HaRxL106 suppresses plant immunity by binding to RADICAL-INDUCED CELL DEATH1. New Phytologist, 220(1), 232–248.

