## Supplemental Data: Suppl. Figures, Materials and Methods, Tables and References for "An oomycete effector that induces shade avoidance like growth and suppresses plant defenses targets the AUX/IAA protein IAA11"

Supplemental Figures
Supplemental Materials and Methods
Supplemental Tables
Supporting References

**Supplemental Figures**

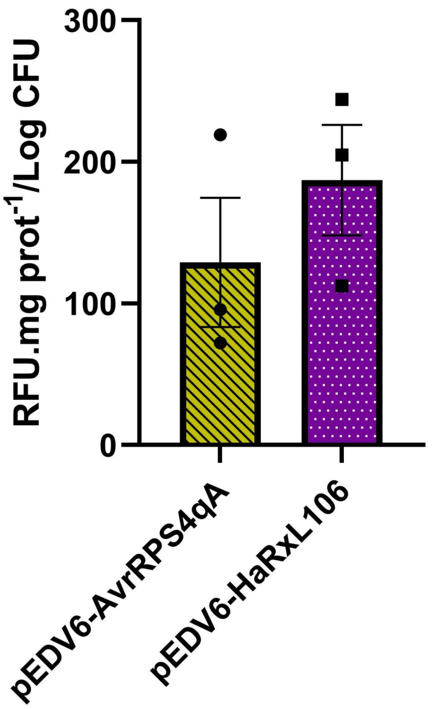


**Supplemental Figure 1: HaRxL106 emerges as a possible auxin-inducing effector.** We used the heterologous Effector Detector Vector (pEDV) system to deliver HaRxL106 via the *P. syringae CUCPB5500* strain into the leaves of *A. thaliana* DR5:GUS plants. Photon emissions (GFP fluorescence) were acquired in 1 square cm of leaves 48 h after inoculation and expressed as relative fluorescence units (RFUs) per mg of protein per Log of CFUs/mL. n=4 plants, 4 leaves per plant. pEDV6-AvrRPS5-AAAA is a negative control.

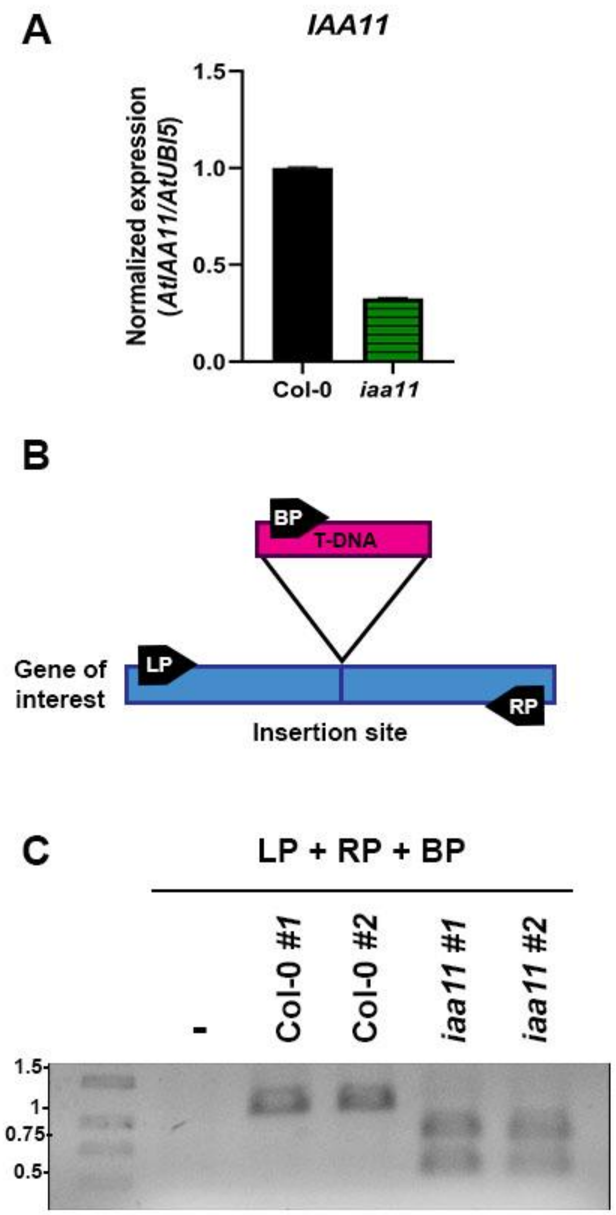


**Supplemental Figure 2: SALK_033787 plant line is an IAA11 knockdown mutant.**
**A**. Expression of IAA11 transcript in Col-0 and *iaa11* 7-day-old seedlings normalized to the reference gene (*AtUBI5*).
**B**. Diagram illustrating the areas where primer hybridization for insertion check is expected. LP stands for left primer, RP for right primer, and BP for primer on the T-DNA. All primers are listed in Supplementary Table 2.
**C**. Electrophoretic run on 1,5% agarose gel of PCRs carried out with the combination of the three primers on the genomic DNA of Col-0 and plants of the line SALK_033787. The absence of band (>1Kb) corresponding to the wild type gene can be observed in lines *iaa11* #1 and 2, as well as the presence of the T-DNA insertion (near 0.75 Kb).

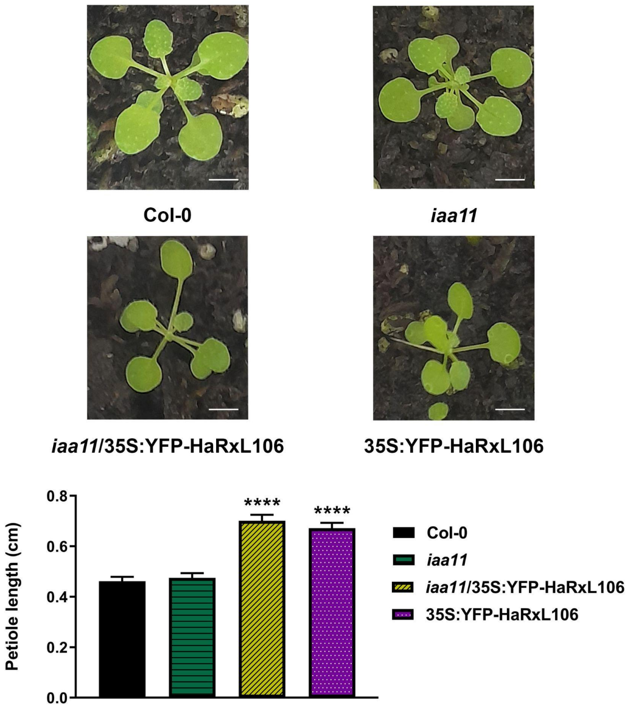


**Supplemental Figure 3: Representative photos and quantification of petiole length of 2-week-old plants.** Statistical significant differences with $P < 0.0001$ according to a one way ANOVA (Tukey's post hoc test) are indicated by asterisks. Scale bars: 5 mm.

**Supplemental Materials and Methods:**

### 1. Plant material and growth conditions

*Arabidopsis thaliana* ecotype Col-0 was used as wild type in this work. Most transgenic or mutant lines in this background were described previously (Supplemental Table 1). *iaa11* mutant (line SALK_033787) was obtained from the NASC/ABRC. This line is reported as a knockdown named *iaa11-4* in Mielecki *et al*., (2022), nevertheless, the insertion in the 5' UTR of the gene might generate a truncated transcript (Supplemental figure 2A). We checked the presence of the T-DNA insertion using 3 different primers: LP: left primer, RP: right primer, BP: primer on the T-DNA (Supplemental Figure 2B, 2C; Supplemental Table 2). Other *iaa11* mutant line requested (SAIL_567-H06.1) is intronic, therefore we decided not to use it for experiments. *iaa11*/35S:YFP-HaRxL106 lines were obtained by transformation by floral dip (Clough and Bent, 1998), with *Agrobacterium tumefaciens* GV3101 containing the 35S:YFP-HaRxL106 expression vector (Supplemental Table 3). Transformants were selected by spraying with a 1/1000 Glyphosate solution (0.2 x$10^3$ mg/mL, Liberty) over three generations. DR5:GUS/pER8:HA-HaRxL106 plants were generated by floral dip, and selected by PCR using the genomic DNA of c.a. 450 T1 plants and specific primers (Supplemental Table 2).

*A. thaliana* or *N. benthamiana* plants were grown on commercial substrate (Grow Mix Multipro, Terrafertil SA). Seeds were stratified for 2 days, then sown directly in soil and maintained at 22 °C, 12:12 photoperiod, 70% humidity and 7000 lux in a growth cabinet (Demetra 380 L, J3 Desarrollos, Mar del Plata, Argentina). For experiments involving Hpa infections, the cabinet temperature was lowered at 18 °C.

For estradiol treatments, seedlings were grown in plates. Seeds were sterilized in Triton X-100/Ethanol as previously described (Bogino *et al.*, 2024), sown in 0.5x Murashige-Skoog medium (MS), 1% agar plates, pH = 5.7, stratified at 4 degrees for 2 days and placed in growth cabinet under the same conditions. HaRxL106 transient expression in pER8:HA-HaRxL106 lines (Bogino *et al.*, 2024) was induced by spraying 9-day old seedlings with 1.25 µM β-estradiol. Seedlings were harvested 24 h post estradiol application, and stained for detection of β-Glucuronidase (GUS) activity as described below.

### 2. Detection of β-Glucuronidase (GUS) activity

The assays with DR5:GUS reporter were conducted on seedlings according to the protocol proposed by Jefferson et al. (1987), performing the infiltration of the staining solution with vacuum and acetone as suggested by Dedow et al. (2022). Seedlings were mounted in 50% glycerol and images were obtained using a camera associated with a magnifying glass (Optika Microscopes Italy). GUS staining intensity was quantitated using the ImageJ software (NIH-USA) with an *ad hoc* macro as described: The RGB images were converted to 8-bit, inverted, the background was subtracted, and the average pixel intensity corresponding to the blue channel in the cotyledon area was determined.

### 3. Screening of effectors with auxin-inducing activity

We used the heterologous Effector Detector Vector (pEDV) system as previously reported (Sohn et al., 2007, Fabro et al., 2011) to deliver c.a. 60 candidate Hpa effectors into the plant cells via the type three secretion system of the *Pseudomonas syringae CUCPB5500* bacterial strain. This bacterial strain lacks 18 effector genes of *P. syringae pv* tomato DC3000 that might interfere with the function of Hpa effectors (Kvitko et al., 2009). HaRxL106 was cloned into pEDV6 vector and the correct in-frame construct was introduced by conjugation into *P. syringae CUCPB5500* strain. Bacterial suspensions at $10^6$ colony forming units per milliliter -CFU/mL- were syringe inoculated into the leaves of Col-0 DR5:GFP reporter plants. After 48 h, 16 leaves of 4 plants were imaged for GFP fluorescence emission (1 square cm per leaf) with a high resolution photon counting system (Photek Ltd, UK). Counts of different leaves (4) from the same plant were averaged and expressed as RFUs (relative fluorescence units). Immediately, imaged areas (1 $cm^2$) were collected, ground in 10 mM $MgCl_2$ pH 5.7 and used to determine bacterial count via serial dilutions and plating in solid LB media supplemented with appropriate antibiotics. An aliquot of the same macerated was used to determine total protein content via Bradford assay. Fluorescence was expressed as RLUs per mg of protein per Log of CFU/mL. Leaves inoculated with *P. syringae* pEDV6 that delivered a truncated version of the bacterial effector AvrRPS4 which is unable to induce DR5:GFP were used as negative controls.

### 4. Yeast-two-hybrid

The ProQuestTM Two-Hybrid System (Invitrogen) was used to perform the Y2H assay. pENTR4-IAA11 containing IAA11 CDS (splicing variant 1, AT4G28640.1, primers used to clone this CDS are listed in Supplemental Table 2) was recombined into the pDEST22 prey vector via Gateway LR reaction. HaRxL106 bait vectors were already published (Bogino *et al.* 2024; Supplemental Table 3). We employed the vectors pDEST32-Krev1 and pDEST22-RalGDS as positive controls in accordance with the manufacturer's guidelines. Negative controls were developed for recording autoactivation by co-transforming yeasts with either one interactor and one empty vector, or both empty vectors. The yeast strain MaV203 was transformed with each combination of plasmids as indicated in Figure 1D, and positive clones were subsequently selected in a Tryptophan-Leucine-lacking commercial medium (SC-WL, Synthetic Defined Yeast Media, MP Biomedicals$^{TM}$). They were then plated on media also lacking Histidine (SC-WLH) and containing 25 mM of 3-Aminotriazole (3-AT) to identify the clones expressing interacting proteins. After five days of incubation at 28 to 30°C, the plates were imaged.

### 5. Bimolecular Fluorescence Complementation Assays

We transiently expressed the YFPn-tagged effector's versions (HaRxL106 FL, HaRxL106-ΔC, and HaRxL106-c58, already published in Wirthmueller *et al.,* 2018 and Bogino *et al*., 2024) with IAA11-YFPc in *Nicotiana benthamiana* leaves. A suspension of 10 mM $MgCl_2$, 10 mM MES buffer pH = 5.7, 150 μM acetosyringone, and *A. tumefaciens* GV3101 expressing the mentioned plasmids was used to infiltrate 1-month-old *N. benthamiana* plants, with $OD_{600}$ = 0.3 of each strain when two proteins were co-expressed; if they were expressed separately, the $OD_{600}$ was 0.4. As a positive interaction control we used the known HaRxL106 target IMPORTIN-α3 -MOS6- (Wirthmueller *et al*., 2015). An *A. tumefaciens* strain containing the P19 silencing suppressor was co-infiltrated at $OD_{600}$ = 0.05 in every instance. Images from confocal microscopy (FV1200, Olympus LS) were obtained 48–72 h after infiltration. Constructs used in these experiments are listed in Supplemental Table 3. The IAA-YFPc expressing clones were generated by recombination of pENTR4-IAA11 no stop into pAM-PAT-35S-YFPc plasmid (Lefebvre et al., 2010) using Gibson Cloning (New England Biolabs) and Gateway Technologies (Thermo Fisher Scientific) as described in Bogino *et al*., 2024.

### 6. Infection assays with Hpa and Pst

7/10-day-old Arabidopsis seedlings were infected with *Hyaloperonospora arabidopsidis* isolate NoCo2 at a concentration of 1–5x$10^5$ conidiospores/mL. Conidiospore number was assessed between 7 - 10 days post inoculation. Seedlings were collected, carefully placed in eppendorf tubes, weighted and vortexed with 1 mL distilled water and 5 μLaliquots used to count conidia with a Neubauer chamber. Conidiospore number per mL was related to the fresh weight (mg) of the infected seedlings. These experiments were repeated 3 times.

For *Pseudomonas syringae pv* tomato DC3000 (Pst) infections, we used 6/8-week-old plants. Freshly Pst grown in solid LB media with rifampicin (100 μg/mL) and kanamycin (50 μg/mL) (2 days, RT) was resuspended in 10 mM $MgCl_2$ to a concentration of 5x$10^5$-$10^6$ CFU/mL and infiltrated with a needless syringe onto the abaxial side of the leaves. After 1 and 3 days, two discs were cut from each of six infected leaves, and treated subsequently as one sample (having n=6 per genotype for each time point). They were ground in 1mL 10 mM $MgCl_2$, serially diluted and plated in LB Agar plates with 50 μg/mL Kanamycin and 100 μg/mL Rifampicin. Colony forming units were counted 48–72 h post incubation. These experiments were conducted three times.

**7. Callose and ROS determinations**

To evaluate callose depositions, leaves of adult plants were detached 24 h post-infection with Pst *hrcC-* mutant strain, chlorophyll was cleared in 96% ethanol, leaves rehydrated in water, and incubated for 1 h in a solution of 0.01 % w/v Aniline Blue (in 150 mM phosphate buffer, pH = 9.5). Leaves were mounted in 50 % glycerol and photographed using an Axiovision camera associated with an Axioplan 135 microscope (Zeiss, Germany), using UV light and a blue filter with a 20X objective. The callose deposits were quantified using the "Find Maxima" function with ImageJ software (NIH-USA) and automated with an ad hoc macro.

To assess the production of ROS, we followed the protocol established by Gómez-Gómez et al. (2000): The afternoon before taking the measurement, 24 discs from fully expanded leaves were cut (2 discs per leaf, one on each side of the main vein) using a #1 punch (0.38 $cm^2$) and incubated in distilled water overnight in the dark. The next morning, they were carefully distributed with a paintbrush in white 96-well multi-well plates containing distilled water. At the moment of the measurement, water was exchanged by a solution of luminol 34 µg/mL (SIGMA #123072), peroxidase 20 µg/mL (SIGMA # P6782), and flg22 100 nM (GenScript # RP19986), and the emitted luminescence was immediately measured using a luminometer (Biotek Synergy HT, Winooski, VT, USA).

**8. Chlorophyll quantitation**

0.307 $cm^2$ leaf discs (cork borer #2) were extracted and chlorophyll cleared on 70% ethanol at 80°C for 10 minutes, then centrifuged at 14000 rpm for 5 minutes in order to determine the total chlorophyll content. The method suggested by Vernon (1960) was used to measure the optical density of the supernatants at 654 nm.

**9. Hypocotyl, petiole and insertion angle measurements**

To measure the hypocotyl length, sterile seeds were sown on 0.8 % Agar water in rectangular plates (70 x 50 x 25 mm) stratified in darkness at 4 °C for one week, transferred to the growing chamber (Percival E-41L2, 8-channel Sci-Brite Lights) with white light (42 µmoles/$m^2$s) for one hour at 22 °C, then maintained in darkness covered with aluminium foil for 23 h and finally uncovered and kept in normal light conditions (12 h white light - 12 h darkness) for 6 days as described by Fankhauser and Casal (2004). On the 6th day, the seedlings were pressed against the media to place them horizontally and images were taken using a scanner. In the case of treatment with far red light at the end of the day (EODFR, Mizuno et al., 2015), an additional period of 10 minutes of Far Red Light (40 µmoles/$m^2$s) was added every day after white lamps turned off.

ImageJ was used to assess the petiole length of two-week-old plants. A protractor was used for measuring the angles of insertion to the rosette in adult plants (8-weeks-old). After that, the leaves were removed, and a caliper was used to determine the length of the petioles.

**10. RT-qPCR**

The SDS-LiCl RNA protocol (Verwoerd *et al.*, 1989) was used to purify total RNA, and reverse transcription was carried out as previously mentioned (Cambiagno *et al.*, 2015). qPCR was carried out using specific primers (Supplemental Table 2) and GreenLight qPCR Master Mix (Inbio Highway, Argentina), with the following protocol: 95°C for three minutes, 40–50 cycles of 20 seconds denaturation at 95°C, 30 seconds annealing at 58°C, and 15 seconds extension at 72°C. Supplemental LinRegPCR software (Ramakers *et al.*, 2003) was used to determine the amplification efficiency for each well, having 3 wells as technical replicates. The transcripts relative quantity was then estimated using the average efficiency (E) for each primer pair, as determined by $E^{(Ct\ control\ sample\ -\ Ct\ test)}$, and normalized to the relative quantity of the reference gene: *AtUBI5* (Ubiquitin 5; At3g62250). These experiments were repeated three times with similar results.

**11. Statistical analysis**
InfoStat (Di Rienzo *et al.*, 2000) and GraphPad Software (GraphPad Software Inc.; San Diego, CA, USA) were used for statistical analyses.

**Supplemental Tables:**

**Supplemental Table 1**: Plant material used in this work

| Plant line | Identifier | Source |
|---|---|---|
| *iaa11* single mutant | SALK_033787 | ABRC/NASC |
| *iaa11*/35S:YFP-HaRxL106 | | This work |
| *bim1* single mutant | N654404/SALK_132178C | Chandler *et al.*, 2009 |
| 35S:BIM1-FLAG | | Liang *et al.*, 2018 |
| 35S:YFP-HaRxL106 | | Wirthmueller *et al.*, 2018 |
| *bim1*/35S:YFP-HaRxL106 | | Bogino *et al.*, 2024 |
| DR5:GUS | | Ulmasov *et al.*, 1997 |
| pER8:HA-HaRxL106 | | Bogino *et al.*, 2024 |
| DR5:GUS/pER8:HA-HaRxL106 | | This work |

**Supplemental Table 2:** Primers used in this work

| Identifier | Sequence | Used for: |
|---|---|---|
| IAA11 LP (SALK_033787) | TCAATCCCATAACCATAAAAAGC | Genotypic analysis of T-DNA insertional mutant lines |
| IAA11 RP (SALK_033787) | GAATCCCATGAAGCTGACATG | Genotypic analysis of T-DNA insertional mutant lines |
| LBb1.3 SALK (primer BP) | ATTTTGCCGATTTCGGAAC | Genotypic analysis of T-DNA insertional mutant lines |
| SAUR-AC_exp_fw | TGAGGAGTTTCTTGGGTGCT | qPCR |
| SAUR-AC_exp_rv | TATTGTTAAGCCGCCCATTG | qPCR |
| XTH19_exp_fw | TTCACGATAATCAAGGGAAAC | qPCR |
| XTH19_exp_rv | AAAGATAGAATGTTGTGAC | qPCR |

| | | |
|---|---|---|
| | GG | |
| UBI5_exp_fw | GTGGTGCTAAGAAGAGGAAGA | qPCR |
| UBI5_exp_rv | TCAAGCTTCAACTCCTTCTTT | qPCR |
| IAA11_exp_fw | ATTAGGTTCACGCACACTAGA | qPCR |
| IAA11_exp_rv | GAACCAATAAACATCCCCCAC | qPCR |
| HaRxL106long_fw | TAGTCTTTCCCGGCTCGCTC | Selection of HaRxL106-expressing plants by semi-quantitative PCR. |
| HaRxL106long_rv | AACTCCTCCCGATTCCCATCA | Selection of HaRxL106-expressing plants by semi-quantitative PCR. |
| IAA11 no start FW | CAAAAAAGCAGGCTCCACTGAAGGCGGTTCCGCTAGTG | Cloning of IAA11 splicing variant 1 CDS in pENTR4 for N-terminal tagging (For Y2H) |
| IAA11 v1 stop RV | GCTGGGTCTAGATATCTCATAATATCATCTGAGCTTTACCAGTAG | Cloning of IAA11 splicing variant 1 CDS in pENTR4 for N-terminal tagging (For Y2H) |
| IAA11 from ATG FW | CAAAAAAGCAGGCTCCACATGGAAGGCGGTTCCGCTAG | Cloning of IAA11 splicing variant 1 CDS in pENTR4 for C-terminal tagging (For BiFC) |
| IAA11 v1 no stop RV | gctgggtctagatatctcgagttCAAAGAGAACATATAACTAACTAAAAGAAAAATGG | Cloning of IAA11 splicing variant 1 CDS in pENTR4 for C-terminal tagging (For BiFC) |
| pENTR4_fw | CCCGCCATAAACTGCCAGG | Sequencing. Anneals on pENTR4 backbone. |
| pENTR4_rv | CGTTGAATATGGCTCATAACACCC | Sequencing. Anneals on pENTR4 backbone. |
| IAA11 middle FW | GGGATGGCCACCAATAAGGA | Sequencing. Anneals on the second exon of all IAA11 splicing variants. |
| IAA11 middle RV | GAGCATTCAGATCGATTTTC | Sequencing. Anneals on |

| | | |
|---|---|---|
| | CTTCC | the second exon of all IAA11 splicing variants. |

**Supplemental Table 3**: vectors used in this work

| **Identifier** | **Source** |
|---|---|
| 35S:YFP-HaRxL106 | Wirthmueller *et al.*, 2018 |
| pER8:HA-HaRxL106 | Bogino *et al.*, 2024 |
| pDEST22-IAA11 | This work |
| pDEST32-HaRxL106 FL | Wirthmueller *et al.*, 2018 |
| pDEST32-HaRxL106-ΔC | Wirthmueller *et al.*, 2018 |
| pDEST32-HaRxL106-c58 | Bogino *et al.*, 2024 |
| YFPn-HaRxL106 | Wirthmueller *et al.*, 2015 |
| YFPn-HaRxL106-ΔC | Wirthmueller *et al.*, 2015 |
| YFPn-HaRxL106-c58 | Bogino *et al.*, 2024 |
| IAA11-YFPc | This work |
| MOS6-YFPc | Wirthmueller *et al.*, 2015 |
| pENTR4-IAA11 | This work |
| pAM-PAT-35S-YFPc | Lefebvre *et al.*, 2010 |

**Supporting references:**

**Bogino, M. F., Lapegna Senz, J. M., Kourdova, L. T., Tamagnone, N., Romanowski, A., Wirthmueller, L., & Fabro, G.** (2024). Downy mildew effector HaRxL106 interacts with the transcription factor BIM1 altering plant growth, BR signaling and susceptibility to pathogens. *The Plant Journal.*

**Cambiagno, D. A., Lonez, C., Ruysschaert, J. M., & Alvarez, M. E.** (2015). The synthetic cationic lipid diC14 activates a sector of the Arabidopsis defence network requiring endogenous signalling components. *Molecular plant pathology*, 16(9), 963-972.

**Chandler, J. W., Cole, M., Flier, A., & Werr, W.** (2009). BIM1, a bHLH protein involved in brassinosteroid signalling, controls Arabidopsis embryonic patterning via interaction with DORNRÖSCHEN and DORNRÖSCHEN-LIKE. *Plant Molecular Biology*, 69, 57-68.

**Clough, S. J., & Bent, A. F.** (1998). Floral dip: a simplified method for Agrobacterium‐mediated transformation of Arabidopsis thaliana. *The plant journal*, 16(6), 735-743.

**Dedow, L. K., Oren, E., & Braybrook, S. A.** (2022). Fake news blues: A GUS staining protocol to reduce false-negative data. *Plant Direct*, 6(2), 1–8.

**Di Rienzo, J., Robledo, W., Casanoves, F., & Balzarini, M.** (2000). InfoStat, Software estadıstico. Córdoba: *Facultad de Ciencias Agropecuarias, Universidad Nacional de Córdoba*.

**Fabro, G., Steinbrenner, J., Coates, M., Ishaque, N., Baxter, L., Studholme, D. J., ... & Jones, J. D.** (2011). Multiple candidate effectors from the oomycete pathogen *Hyaloperonospora arabidopsidis* suppress host plant immunity. *PLoS pathogens*, 7(11), e1002348.
**Fankhauser, C., & Casal, J. J.** (2004). Phenotypic characterization of a photomorphogenic mutant. *The Plant Journal*, 39(5), 747-760.
**Gómez-Gómez L, Boller T.** (2000) FLS2: an LRR receptor-like kinase involved in the perception of the bacterial elicitor flagellin in Arabidopsis. Mol Cell. Jun;5(6):1003-11.
**Jefferson, R. A., Kavanagh, T. A., & Bevan, M. W.** (1987). GUS fusions: beta-glucuronidase as a sensitive and versatile gene fusion marker in higher plants. *The EMBO journal*, 6(13), 3901-3907.
**Kvitko, B. H., Park, D. H., Velásquez, A. C., Wei, C. F., Russell, A. B., Martin, G. B., ... & Collmer, A.** (2009). Deletions in the repertoire of Pseudomonas syringae pv. tomato DC3000 type III secretion effector genes reveal functional overlap among effectors. *PLoS pathogens*, 5(4), e1000388.
**Lefebvre, B., Timmers, T., Mbengue, M., Moreau, S., Hervé, C., Tóth, K., ... & Ott, T.** (2010). A remorin protein interacts with symbiotic receptors and regulates bacterial infection. *Proceedings of the National Academy of Sciences*, 107(5), 2343-2348.
**Liang, T., Mei, S., Shi, C., Yang, Y., Peng, Y., Ma, L., ... & Liu, H.** (2018). UVR8 interacts with BES1 and BIM1 to regulate transcription and photomorphogenesis in Arabidopsis. *Developmental cell*, 44(4), 512-523.
**Mielecki, J., Gawroński, P., & Karpiński, S.** (2022). Aux/IAA11 is required for UV-AB tolerance and auxin sensing in Arabidopsis thaliana. *International Journal of Molecular Sciences*, 23(21), 13386.
**Ramakers, C., Ruijter, J. M., Lekanne Deprez, R. H., & Moorman, A. F. M**. (2003). Assumption-free analysis of quantitative real-time polymerase chain reaction (PCR) data. *Neuroscience Letters*, 339(1), 62–66. https://doi.org/10.1016/S0304-3940(02)01423-4
**Sohn, K. H., Lei, R., Nemri, A., & Jones, J. D.** (2007). The downy mildew effector proteins ATR1 and ATR13 promote disease susceptibility in Arabidopsis thaliana. *The Plant Cell*, 19(12), 4077-4090.
**Ulmasov, T., Murfett, J., Hagen, G., & Guilfoyle, T. J.** (1997). Creation of a Highly Active Synthetic AuxRE. *Society*, 9(November), 1963–1971.
**Vernon, L. P.** (1960). Spectrophotometric Determination of Chlorophylls and Pheophytins in Plant Extracts-Corrections. *Analytical Chemistry*, 32(11), 1414.
**Verwoerd, T. C., Dekker, B. M., & Hoekema, A.** (1989). A small-scale procedure for the rapid isolation of plant RNAs. *Nucleic acids research*, 17(6), 2362.
**Wirthmueller, L., Roth, C., Fabro, G., Caillaud, M. C., Rallapalli, G., Asai, S., ... Jones, J.D. & Banfield, M. J.** (2015). Probing formation of cargo/importin-α transport complexes in plant cells using a pathogen effector. *The Plant Journal*, 81(1), 40-52.
**Wirthmueller, L., Asai, S., Rallapalli, G., Sklenar, J., Fabro, G., Kim, D. S., Lintermann, R., Jaspers, P., Wrzaczek, M., Kangasjärvi, J., MacLean, D., Menke, F. L. H., Banfield, M. J. & Jones, J. D.** (2018). Arabidopsis downy mildew effector HaRxL106 suppresses plant immunity by binding to RADICAL-INDUCED CELL DEATH1. *New Phytologist*, 220(1), 232-248.
